# Combined CRISPRi and proteomics screening reveal a cohesin-CTCF-bound allele contributing to increased expression of *RUVBL1* and prostate cancer progression

**DOI:** 10.1101/2023.01.18.524405

**Authors:** Yijun Tian, Dandan Dong, Zixian Wang, Lang Wu, Jong Y Park, Gong-Hong Wei, Liang Wang, PRACTICAL/ELLIPSE consortium

**Author notes:** ^6^These authors contributed equally (Yijun Tian, Dandan Dong). ^7^Check supplementary information for the full author list (PRACTICAL/ELLIPSE consortium).

## Abstract

Genome-wide association studies along with expression quantitative trait loci (eQTL) mapping have identified hundreds of single nucleotide polymorphisms (SNPs) and their target genes in prostate cancer (PCa), yet functional characterization of these risk loci remains challenging. To screen for potential regulatory SNPs, we designed a CRISPRi library containing 9133 guide RNAs (gRNAs) to target 2,166 candidate SNP sites implicated in PCa and identified 117 SNPs that could regulate 90 genes for PCa cell growth advantage. Among these, rs60464856 was covered by multiple gRNAs significantly depleted in the screening (FDR<0.05). Pooled SNP association analysis in the PRACTICAL and FinnGen cohorts showed significantly higher PCa risk for the rs60464856 G allele (pvalue=1.2E-16 and 3.2E-7). Subsequent eQTL analysis revealed that the G allele is associated with increased *RUVBL1* expression in multiple datasets. Further CRISPRi and xCas9 base editing proved the rs60464856 G allele leading to an elevated *RUVBL1* expression. Furthermore, SILAC-based proteomic analysis demonstrated allelic binding of cohesin subunits at the rs60464856 region, where HiC dataset showed consistent chromatin interactions in prostate cell lines. *RUVBL1* depletion inhibited PCa cell proliferation and tumor growth in xenograft mouse model. Gene set enrichment analysis suggested an association of *RUVBL1* expression with cell-cycle-related pathways. An increased expression of *RUVBL1* and activations of cell-cycle pathways were correlated with poor PCa survival in TCGA datasets. Together, our CRISPRi screening prioritized about one hundred regulatory SNPs essential for prostate cell proliferation. In combination with proteomics and functional studies, we characterized the mechanistic role of rs60464856 and *RUVBL1* in PCa progression.

## Introduction

Among all cancer types, prostate cancer (PCa) accounted for 26% of 970,250 new cancer cases and caused 11% of 319,420 cancer-related deaths in US males in 2021. ^1^ As a cancer type with strong genetic predispositions, PCa has been extensively investigated in genome-wide association studies (GWAS) ^2^ to determine susceptible variants associated with increased disease risk and aggressiveness. ^2,3^ It has been reported that the joint contribution of GWAS identified loci to PCa risk is nearly 20%.^4^ Although GWAS have been highly productive, only a few risk loci have been functionally characterized. ^5–8^ Thus far, as these risk SNPs are found in non-coding parts of the genome, it is believed that many of them (or their closely linked SNPs) would alter regulatory element activities and quantitatively change gene expression rather than directly mutating protein sequences. ^9–12^ With tremendous large-scale GWAS findings, especially highly reproducible ones, it is believed that some non-coding variants play a subtle but profound role in PCa initiation, progression, and metastasis by modulating the expression of certain susceptible genes.

To dissect functional variants and their target genes in PCa, bioinformatic and benchtop approaches are applied to prioritize and validate the causal variants. ^13–15^ Curated databases such as ChIP-atlas, ^16^ ENCODE, ^17^ JASPAR, ^18^ and GTEx^19^ provided abundant resources to estimate the genetic contribution of GWAS variants to PCa and other cancer susceptibilities. However, large-scale genetic assays are often needed to interrogate endogenous variant loci and directly characterize consequent phenotype changes to determine the biological effect. ^20–22^ One such assay is lentiviral-based Cas9 screening, which has emerged as a powerful tool for evaluating the biological significance of certain genes on a large scale. ^23,24^ Compared to canonical wildtype Cas9 based screening, dead Cas9 (dCas9) forms steric hindrance according to the gRNA sequence and induces transcription repression if fused to repressor peptides KRAB (Krüppel-associated box). The dCas9-KRAB has been proven to specifically decrease target gene expression without strand cleavage and is used for screening regulatory elements in mammalian cells. ^25,26^

Most SNPs are believed to function as regulatory switches in the human genome. With the eliminated nuclease activity, ^26^ we hypothesized that dCas9-based CRISPRi could be used to interfere with regulatory sequences at SNP loci and faithfully mimick the transcription alteration caused by single base differences. To test this hypothesis, we first established multiple stable dCas9-KRAB prostate cell lines and designed an unbiased, highly reproducible gRNA library that targeted candidate SNPs at PCa risk loci. We then performed negative selection for potential SNPs conferring growth advantage. We finally provided a detailed analysis of an SNP-gene pair for its functional role using prostate cell lines and mouse model and performed a successful proteomics identification of transcriptional regulators in transforming the gene regulatory effects. Our results support the CRISPRi-based approach at risk loci for regulatory SNP screening.

### Method

#### eQTL-based SNP selection at prostate cancer risk loci

The rationale for the regulatory screening was explained in **Supplementary Figure 1A** and **1B**. In brief, to select candidate SNPs, we first retrieved cis-eQTL data from our benign prostate tissues^27^ and identified all SNPs with gene-wise FDR≤1E-3. We then applied ENCODE annotations (including histone modification, common transcription factor binding, and DNase hypersensitivity in prostate cell lines) to filter for candidate functional SNPs.

#### gRNA selection and library pool amplification for candidate SNPs

To design gRNAs for the candidate SNPs, we retrieved DNA sequences surrounding each SNP (± 23bp). We used the CRISPOR program (http://crispor.tefor.net/) for in-silico selection. We assembled the gRNA oligo pool into the lentiGuide-Puro (Addgene #52963) backbone and transformed the pooled oligos into highly sensitive Endura ElectroCompetent Cells (Lucigen) to generate plasmid libraries.

#### dCas9/KRAB stable cell lines and gRNA library processing

To establish stable cell lines, we packaged lenti-dCas9-KRAB-blast (Addgene #89567) into lentiviral particles with HEK293FT cells and used ten-fold concentrated virus particles to transduce cell lines. After blasticidin selection, the stable dCas9 expressions were verified with western blots. Meanwhile, the gRNA virus library (packaged from lentiGuide-Puro) was titered in each dCas9 stable cell line. To achieve low MOI infection, we optimized the cell number and virus amount over 72-hour puromycin selection, such that the non-infected group would be eliminated by 95% to 99%, while the library group retained 30% to 40% variability. The cell viability was measured with the CellTiter-Glo® Luminescent kit (Promega G7570). After confirming low MOI transducing, we removed puromycin from the medium and continued the cell culture for 21 days. We isolated genomic DNA at baseline D1 (day 1) and endpoint D21 (day 21).

#### gRNA readout sequencing

We used the Illumina HiSeq platform to sequence the gRNA readout amplicons. We aimed for at least a 500-fold library size depth for each replicate to ensure quantification accuracy.

#### Data QC and analysis

To quantify the representation of each gRNA, we used a python script “count_spacer.py” developed by Feng Zhang lab to scan the FASTQ file for perfectly matched hits and generate raw read counts for each experiment. We then used principal components analysis (PCA) to evaluate the similarity of the plasmid library, baseline, and screening endpoint gRNA representation. To determine SNP alleles conferring growth advantage in these cell lines, we used RIGER (RNAi gene enrichment ranking) extension (https://software.broadinstitute.org/GENE-E/extensions.html) to calculate a rank list for SNPs or alleles with the most depleted gRNA representation. ^28^ RIGER program ranks gRNAs according to their depletion effects and then identifies the SNP targeted by the shRNAs.

#### Plasmid construction and siRNA design

To enable the fluorescence-based cell sorting, copGFP ORF was assembled into base editor plasmid^29^ xCas9(3.7)-ABE(7.10) (Addgene #108382) with NEBuilder® HiFi DNA Assembly Master Mix (New England Biolabs). A 223-bp flanking sequencing surrounding rs60464856 was amplified from RWPE-1 cells (rs60464856 heterozygote) and further subcloned into the pGL3-basic vector between NheI and XhoI sites using NEBuilder® HiFi DNA Assembly Master Mix. Small hairpin RNA (shRNA) targeting *RUVBL1*, primers, and gRNA gblock sequences were listed in **Supplementary Table S1**.

#### CRISPR base editing

To change the rs60464856 A allele to G allele, we created a GFP labeled xCas9(3.7)-ABE(7.10) plasmid based on the backbone from David Liu’s lab. ^29^ gRNA template was synthesized in a gblock fragment with a hU6 promoter and a scaffold (https://benchling.com/protocols/10T3UWFo/detailed-gblocks-based-crispr-protocol). Since rs60464856 is located in the base editing window, the adenine base editor can be directed to the SNP site and catalyze A into the G allele. We co-transfected 2.5 ug xCas9(3.7)-ABE7.10 plasmid and 1.2 ug gRNA gblock into cells 80% confluent in each well of the 6-well plate. After 48 hours, GFP-positive cells were sorted by flow cytometry and collected for allele dosage quantification. The GFP-positive cells were seeded into single clones once editing efficiencies were above 5%. After expanding for ten days, we used amplification-refractory mutation system (ARMS) PCR to genotype direct lysate from every clone. We also used Sanger sequencing to verify the germline change of each single clone.

#### Real-time PCR and Western blot analysis

Total RNA was extracted from cells using the Direct-zol RNA Miniprep Kit (Zymo Research). One microgram of total RNA was reverse transcribed by iScript cDNA Synthesis Kit (BioRad). Quantification reactions were performed with PowerUp SYBR Green Master Mix (Thermo Fisher Scientific) on the CFX96 Touch real-time PCR system (BioRad). The primers were listed in **Supplementary Table S1**. Total protein was extracted and electrophoresed as described previously, ^30^ with minor modification using Tricine Sample Buffer and Mini-PROTEAN® Precast Gels (BioRad). SuperSignal West Pico Chemiluminescent Substrate (Thermo Fisher Scientific) produced luminescent signals on the LICOR imaging system. Captured images were aligned in Photoshop and assembled in Illustrator.

#### Luciferase reporter assay

The cells were seeded into a 24-well plate. After 12 to 16 hours, 500 ng of pGL3 reporter plasmids were transfected to each well using Lipofectamine 3000. The media were replaced after transfection for 24 hours. After 48 hours of transfection, the cells were lysed for the luciferase assay according to Dual-Luciferase® Reporter Assay (E1960, Promega) protocol. The luminescence signals were measured with the GlowMax plate reader. After normalized to Renilla luciferase readout, relative firefly luciferase activities driven by corresponding promoters were represented by luminescence unit fold changes.

#### Allele-specific proteomics screening with stable isotope labeling by amino acid in cell culture (SILAC)

The BPH1, DU145, and PC3 cells were grown in SILAC RPMI 1640 medium (ThermoFisher 88365) for five passages before harvesting for nuclear protein extraction (ActiveMotif 40010). After confirming that heavy amino acid labeling efficiency reached 99.9%, the nuclear extracts were applied to the desalting spin column (ThermoFisher 89882) to remove excessive ions. The DNA baits harboring rs60464856 A and G alleles were produced according to a previous publication. ^31^ For each binding reaction, 2ug of purified DNA baits were conjugated to 25 uL Streptavidin Dynabeads (ThermoFisher 65001). The clean conjugated beads were incubated with 12.5 uL precleared nuclear protein at 4 degrees overnight. The incubated beads were washed five times and combined for two parallel quantitative MS runs to get the allelic protein binding ratio. Qualified proteins with allelic binding were defined as **a)** concordant allele ratio changes: log2(A/G)×log2(G/A)<0; **b)** drastic allele ratio differences: |log2(A/G)-log2(G/A)|≥2. For proteins with allelic hits, we further narrowed down to those with known DNA binding functions according to UniProt databases (https://www.uniprot.org/).

#### Chromatin immunoprecipitation (ChIP) qPCR assays

Since the rs60464856 locus was demonstrated to reside in an insulation region in an analysis of previously published Hi-C data, we adapted our ChIP qPCR protocols for chromosome conformation capturing. ^32^ Compared with conventional ChIP assay protocol, our protocol applied dual crosslinking to maximally preserve chromatin contacts. ^33^ Each ChIPed DNA sample was tested in four qPCR reactions, including **a)** rs60464856 locus enrichment primer pair; **b)** rs60464856 upstream locus enrichment primer pair; **c)** rs60464856 A allele-specific primer pair; **d)** rs60464856 G allele-specific primer pair.

#### In vivo xenograft mouse model

Animal experiments were performed according to the protocol approved by the Institutional Animal Care and Use Committee of Fudan University. Nude mice (6 weeks males) were purchased from GemPharmatech Company (Jiangsu, China) and maintained in a pathogen-free environment. PC3 was grown in RPMI1640 containing 10% FBS, 100 U/mL penicillin, and 100 μg/mL streptomycin in a humidified CO_2_ incubator. Before injection into the mice, the cells were harvested by trypsinization and washed two times with PBS. And then PC3 cells were resuspended in 100 μL serum-free medium, mixed with 50% Matrigel (BD Biosciences) and injected (5 × 10^6^/site) subcutaneously into hind flank of each mouse. Tumor volume was measured using digital caliper once a week and calculated using the formula: V = (L×W^2^)/2 (L, length; W, width; all parameters in millimeters). After 4 weeks, the mice were sacrificed, and tumors were taken for weight measurement.

## Results

### Selected candidate SNPs, gRNAs and transfected libraries

We first screened ENCODE database and identified 2664 SNPs (TARGET) with strong epigenomic signals (**Supplementary Figure 1C**). We then applied a Bayesian framework using summary statistics to calculate a posterior inclusion probability (PIP) to predict SNP functionality. We included 5 SNPs with the highest PIP score for each eQTL-associated gene as another PIP SNP category (N = 194). Finally, we included 641 control SNPs (CTL) that showed strong eQTL signals but without epigenomic features. After removing duplicated SNPs, we selected 3408 SNPs for gRNA design. We scanned these SNPs with the CRISPOR program and eventually designed 9133 gRNAs, including 100 control sequences that did not target any genome loci (NCG) (**Supplementary Figure 1D**). The exact sequence design can be found in **Supplementary Table S2**.

To confirm the stable dCas9 expression, we used western blots to confirm the stable expression of dCas9 after one month of transduction (**Supplementary Figure 2A**). We ensured low MOI infection by measuring gRNA lentiviral library function titer in 72 hours (**Supplementary Figure 2B**) and visualized a 261bp gRNA amplicon for each sample on agarose gel for quantification by high-throughput sequencing (**Supplementary Figure 2C**). The sequencing summary for each sample is shown in **Supplementary Figure 2D**. We also visualized the normalized count in each replicate endpoint and observed high correlations in all three cell lines (**Supplementary Figure 2E-2G**).

### CRISPRi screening identified top candidates of regulatory SNPs

We first performed PCA analysis and found tight clustering between baseline and plasmid libraries (**Supplementary Figure 3A**), suggesting faithful representations of original gRNA libraries in transfected cell lines. This analysis also found highly diverse but cell line-dependent distribution in the end libraries, indicating gRNA profile changes by the selection process. We then normalized the read count using the total perfect count in each sample and calculated fold change by dividing the read count in the end library by the read count in the baseline library. The subsequent RIGER analysis showed 779 gRNAs targeting 117 SNPs with permutation test FDR <0.1 in both replicates (**Figure 1B**). Further analysis did not find significant correlations between fold change and gRNA specificity score (**Supplementary Figure 3B-3D**), suggesting minimal off-target effects of these selected gRNAs. When comparing end and baseline libraries, we found significant gRNA depletion in BPH1, DU145, and PC3 screening experiments (**Figure 1C-1E**). When comparing gRNA fold change between different categories of candidate SNPs, we found significantly higher growth depletion only in BPH1 cells with gRNA targeting the PIP and TARGET SNPs (**Supplementary Figure 3E-3G**). We also found that the SNPs selected as screening candidates had a significantly higher proportion of residing in human gene transcription start sites (**Figure 1F**) and tended to be allelically bound to transcriptional factor binding according to ANANASTRA annotation (**Figure 1G**). ^34^ We plotted the gRNA representation before and after the growth selection and highlighted representative SNPs in each cell line (**Figure 1H-1J**). The raw and normalized gRNA count, and the eQTL mapped with the candidate SNP are listed in **Supplementary Table S2**. The RIGER analysis output is listed in **Supplementary Table S3**.

**Figure 1.**
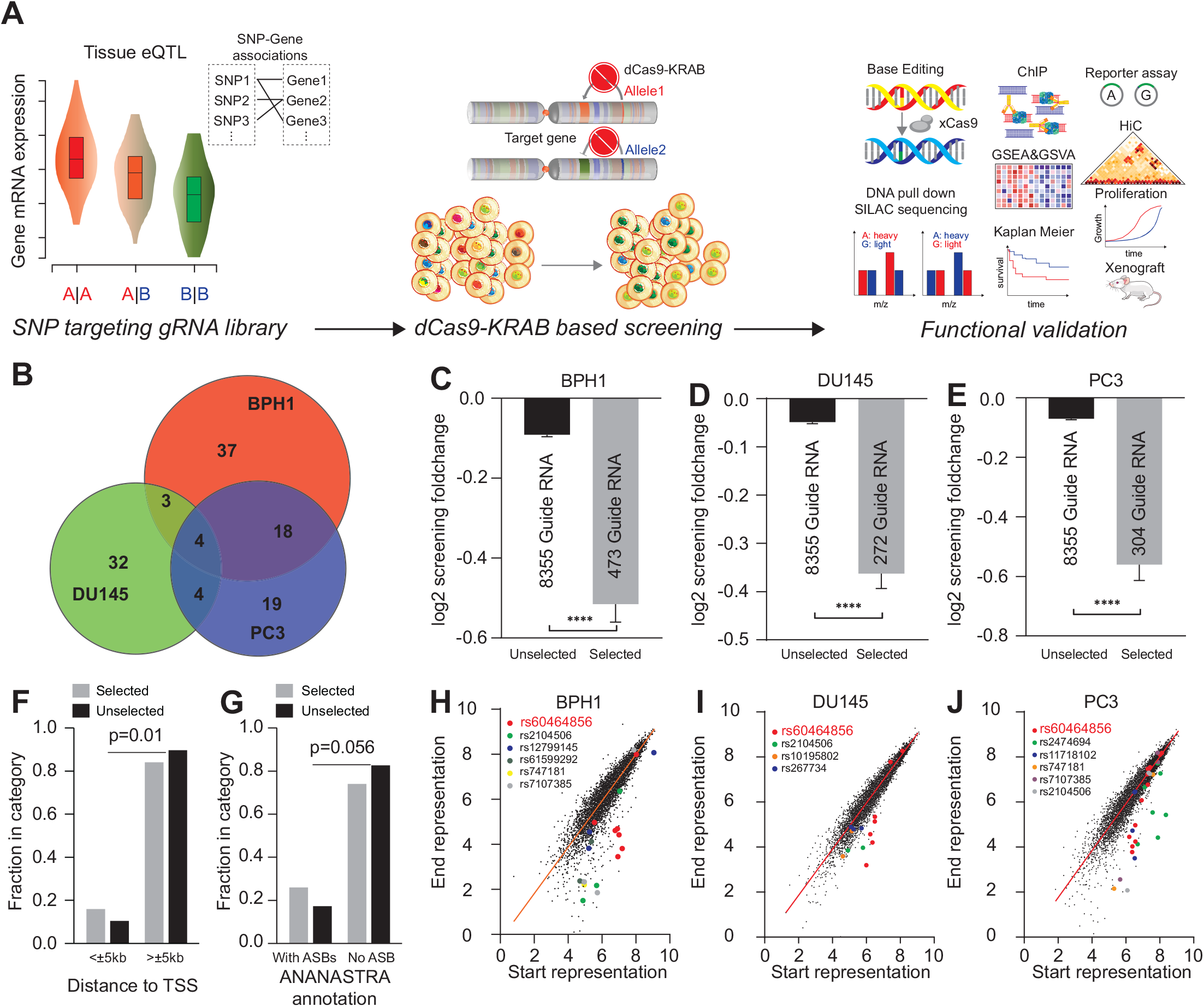
**A.** Study design of the current project, including eQTL target selection, CRISPRi screening, and functional validation. **B.** Overall SNP candidates in BPH1 (red), DU145 (green) and PC3 (blue) cells over 3-week screening. **C-D.** Comparison of the gRNA fold change between the selected and unselected SNPs in BPH1 (**C**), DU145 (**D**) and PC3 (**E**) cells. **F.** Enrichment of the selected SNPs in transcription start sites (TSS) compared to the unselected. **G.** Enrichment of the selected SNPs in allele-specific binding (ASB) annotation according to the ANANASTRA database compared to the unselected. **H-J.** Demonstration of gRNA representation for the selected SNPs in BPH1 (**H**), DU145 (**I**) and PC3 (**J**) cells. gRNA targeting the main validation SNP rs60464856 is highlighted in red.

### rs60464856 showed a regulatory role in *RUVBL1* expression underpinning prostate cancer susceptibility

Among the 117 SNPs showing significant growth inhibition over a 3-week screening, the SNP rs60464856 was consistently selected in all tested prostate cell lines. The SNP sequence was targeted by 10 gRNAs for A and 6 gRNAs for G allele. We observed significant A allele depletion (fold change _≤_ 0.75) in 10 gRNAs in BPH1, 5 gRNAs in DU145 and 10 gRNAs in PC3 cells (**Figure 2A**). The rs60464856 is located in a previously identified risk locus, and the G allele was associated with a 10% increased prostate cancer risk in 107,247 cases and 127,006 controls (pvalue = 1.2E-16) (**Figure 2B**). ^35^ Consistently, a phenome wide association analysis (PheWAS) in the FinnGen cohort (n = 342 499) with 2202 endpoints revealed the strongest association of rs60464856 with malignant neoplasm of prostate (11,590 cases and 110,189 controls; pvalue = 3.2E-7) (**Supplementary Figure 4A**). ^36^ We further showed that among the *RUVBL1* eQTL loci, rs60464856 resided in a linkage disequilibrium block with the second-best significance (**Figure 2C**). Additionally, the rs60464856 G allele was significantly associated with elevated *RUVBL1* gene expression in GTEx, Mayo, ^27,37,38^ and TCGA prostate eQTL^39^ datasets (**Figure 2D**). We also used three gRNAs targeting the rs60464856 locus to transiently transfect the dCas9 stable cells and observed a consistent knockdown effect on *RUVBL1* gene expressions in all three prostate cell lines (**Figure 2E**). To evaluate the nuances of endogenous allele transition contributing to downstream gene expression, we applied nickase Cas9 (xCas9) base editing technology, which featured high conversion efficiency and minimal indel rate, to faithfully mutate the rs60464856 locus in prostate cells (**Figure 2F**). Finally, we generated multiple isogenic subclones with accurate base conversion and confirmed significant increase of endogenous *RUVBL1* expression by the G allele in BPH1 and DU145 (**Figure 2G**).

**Figure 2.**
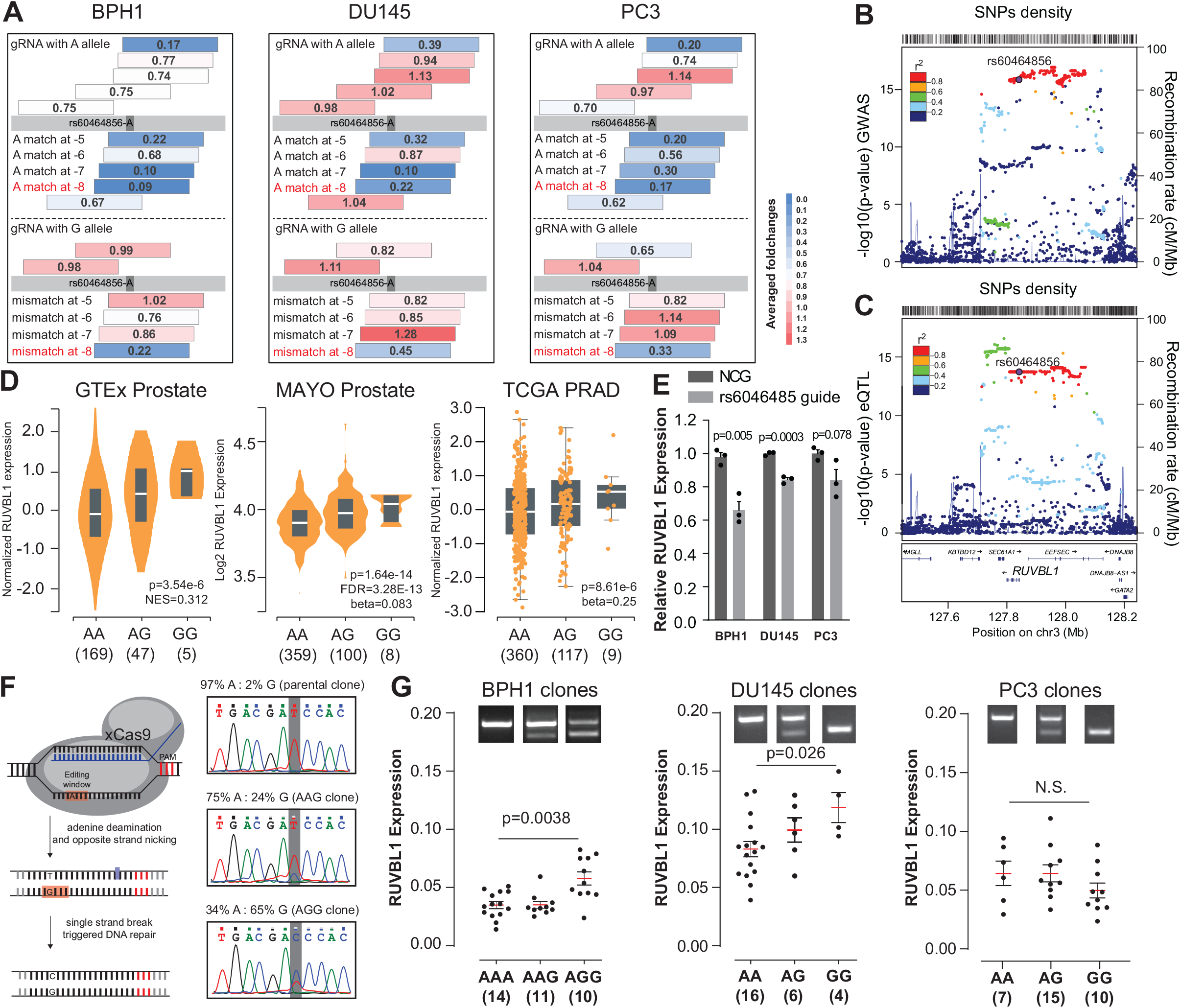
**A.** Screening fold change of rs60464856 gRNA in BPH1, DU145 and PC3 cells. The Grey bar indicates rs60464856 locus in the germline, and colored strips above or below the bar indicate gRNA binding Crick or Watson strands of the germline DNA. Numbers in the center show the average fold change for two screening replicates. **B.** Locus plots for rs60464856 locus GWAS significance in the PRACTICAL cohort. **C.** Locus plots for *RUVBL1* eQTL significance in the MAYO prostate dataset. **D.** *RUVBL1* gene expressions in dCas9-KRAB stable prostate cells transfected with non-targeting control guide (NCG) and rs60464856 targeting gRNAs. The individual point indicated different gRNA clones. **E.** eQTL associations between *RUVBL1* gene expression and rs60464856 genotypes in GTEx, MAYO and TCGA prostate cohorts. **F.** Schematic figure about xCas9-based A to G base editing in prostate cell lines. **G.** *RUVBL1* gene expressions in rs60464856 base edited clones in BPH1, DU145 and PC3 cells.

### rs60464856 bound to cohesion subunits allelically and mediated chromatin interactions

To evaluate potential allelic protein binding at rs60464856, we searched for ChIP-seq data collections, such as cistrome and ChIP-ATLAS, for potential transcription factor bindings. However, we did not find highly convincing evidence showing allelic TF binding on this locus. Since multiple datasets report histone modification signals on the rs60464856 locus, we then performed ChIP-qPCR to evaluate the locus-specific enrichment for active histone modification markers (H3K4me1, H3K4me3, and H3K27ac) in the base-edited subclones. We found distinct histone modification status in PC3 clones, especially for H3K4me1 on the rs60464856 locus (**Figure 3A**). Consistently, we found the enrichment of H3K4me1, H3K4me3, and H3K27ac modifications on rs60464856 in multiple cell lines (**Figure 3B**), suggesting a robust regulatory potential of this locus in the prostate. With a recent HiC dataset, ^40^ we also visualized the long-range interactions surrounding rs60464856 locus. By summing the normalized count from the interaction hot spot on both sides of the rs60464856 locus, we calculated the left-to-right ratio (L/R) in each cell line. This analysis showed that PC3 cells had only 60% interaction compared to DU145 cells (**Figure 3C**). In contrast, the interactions L/R ratios were roughly equal in VCaP and LNCaP cells (**Figure 3D**). To confirm the transcription activity of the rs60464856 locus, we tested its flanking sequence (chr3: 128,123,257-128,123,479, negative strand) using a reporter assay and found higher promoter activity of the G allele than the A allele in these cell lines (**Figure 3E**). We then applied SILAC-based proteomics to detect possible transcription factors or DNA-binding proteins. This assay took advantage of isotype-labeled nuclear extract in DNA pull-down reactions, thus converting the protein binding difference into the ratio of different molecular weights (**Figure 3F**). This proteomics analysis identified several cohesion subunits bonded to the rs60464856 A than the G allele with the BPH1 nuclear extract (**Figure 3G**). To further elucidate whether the cohesion could bind to endogenous rs60464856 locus allelically, we used ChIP assays specific to the rs60464856 locus from BPH1 and PC3 base edited populations. In BPH1 cells, we found significant locus enrichment (**Figure 3H**) for SMC1A, SMC3, and CTCF, and the A allele preference for SMC3 and CTCF (**Figure 3I**). In PC3 cells, we only found minor locus enrichment for SMC3 and CTCF (**Figure 3J**), and no allele preference was observed for all tested antibodies (**Figure 3K**). We also explored the unique allelic binding role of SMC3, using existing ChIP-seq datasets^41^ (GSE49402 and GSE36578) that quantified SMC1A and SMC3 binding in the human genome (**Supplementary Figure 5A**). We performed STRMEME motif scan with the private peak region of SMC1A and SMC3, and only found that only SMC3 private peaks included an outstanding significant motif in both cell lines (**Supplementary Figure 5B).** Through STRMEME -Tomtom comparison, we also found that this motif was highly similar to the CTCF binding site (MA0139.1) (**Supplementary Figure 5C**). More importantly, we found that the rs60464856 A allele is located in the CTCF zinc finger seven interaction domain and is consistently preferred by multiple versions of the CTCF motif. ^42^ This result implicated a potential mechanism about how the rs60464856 protective allele mediated the cohesion-CTCF complex formation and supported our observation in the HiC datasets (**Figure 3C-3D**).

**Figure 3.**
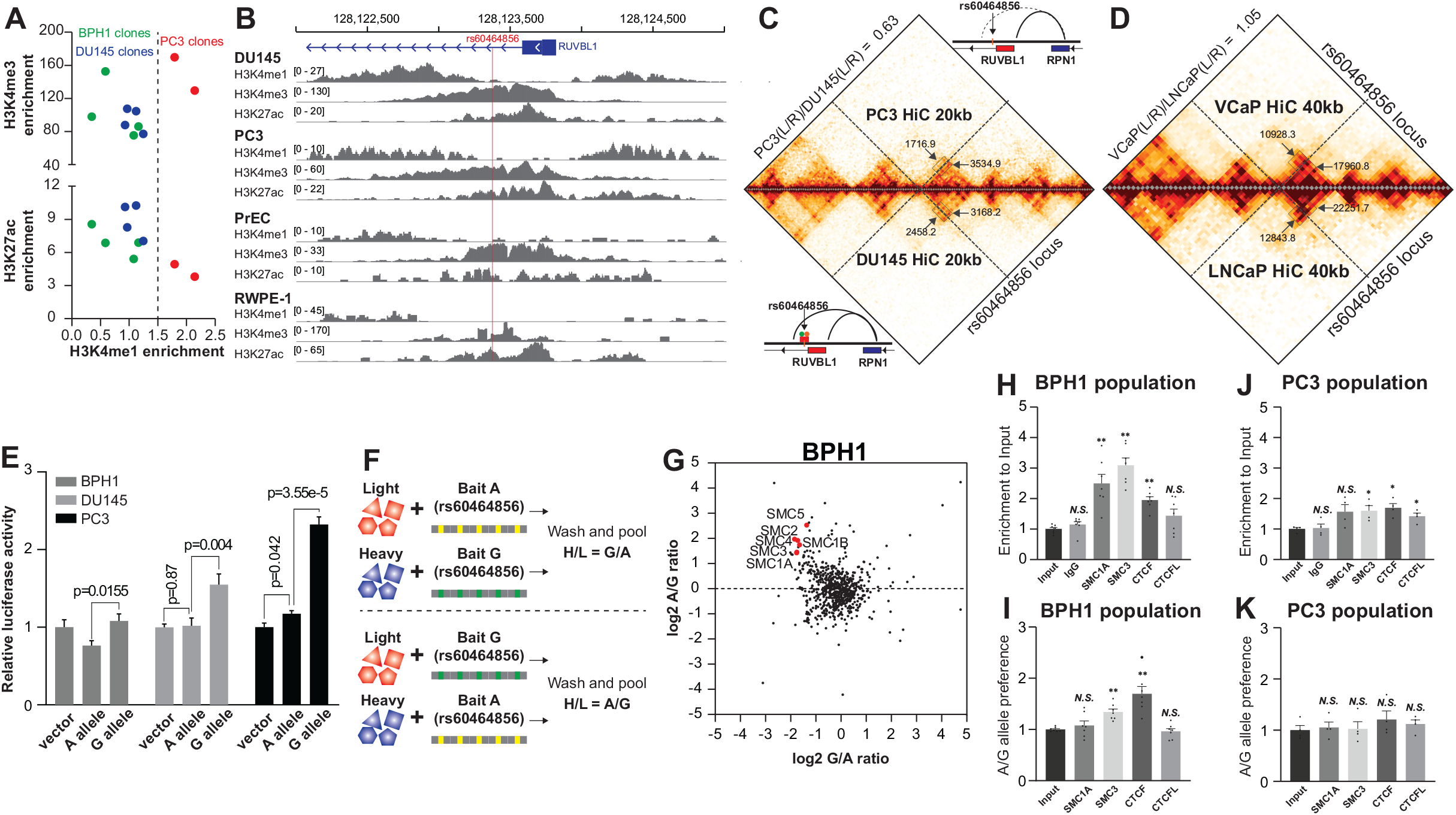
**A.** H3K4me1, H3K4me3 and H3K27ac modifications on rs60464856 locus in BPH1, DU145 and PC3 clones. **B.** ChIP-seq signals of H3K4me1, H3K4me3 and H3K27ac modifications on rs60464856 locus in prostate cells. **C-D.** HiC interaction heatmap in PC3 (C top), DU145 (C bottom), VCaP (D top) and LNCaP (D bottom) cells. L and R indicate interaction ICE normalized read count on the left and right sides of rs60464856 bin. **E.** Luciferase reporter assay comparing rs60464856 allele transcription activities in BPH1, DU145 and PC3 cells.**F.** SILAC-based DNA pull-down proteomics pipeline in identifying allelic protein binding on rs60464856 locus. **G.** Allelic proteomics with BPH1 SILAC extracts. **H-K**. ChIP-qPCR measuring rs60464856 locus enrichment and allele-specific binding to SMC1A, SMC3, CTCF and CTCFL proteins in BPH1 (**H-I**) and PC3 (**J-K**) base edited populations.

### *RUVBL1* knockdown inhibited prostate cell proliferation by downregulating cell cycle related pathways

To evaluate the oncogenic role of the *RUVBL1* gene, we examined the perturbation effect of *RUVBL1* by CRISPR or RNAi screening in DepMap portal. We found that *RUVBL1* was a common essential gene in most human cell lines, with an even better dependency score than the median of all essential genes for the prostate cell lines used in our screening, including BPH1, DU145 and PC3 cells (**Figure 4A**). To further characterize the function of *RUVBL1*, we generated stable cell lines infected with small hairpin RNA (shRNA) lentiviral particles and verified that the RUVBL1 had been successfully knockdown at protein levels (**Figure 4B**). We further monitored the growth of the stable cell lines with daily Incucyte scans and found that *RUVBL1* knockdown by both shRNAs significantly suppressed proliferation in BPH1 (**Figure 4C**), DU145 (**Figure 4D**), and PC3 (**Figure 4E**). We also observed drastic reductions in colony formation for the *RUVBL1* knockdown group in BPH1, DU145, and PC3 cells (**Figure 4F**). To characterize the transcriptome alteration caused by *RUVBL1* downregulation, we quantified RNA profiling of BPH1 and PC3 cells and calculated the GSVA score for HALLMARK geneset collection. Interestingly, multiple cell-cycle-related pathways, including the MYC target, E2F target, and G2M checkpoint genes, showed significant enrichment with *RUVBL1* expression (**Figure 4G-4H**). We further visualized the gene expression changes in these significantly enriched pathways and found consistent trends with the *RUVBL1* expression levels (**Figure 4I**). To characterize the tumorigenic effect of the *RUVBL1* in vivo, we performed xenograft mouse experiments with the PC3 stable cell lines. We found that the *RUVBL1* knockdown significantly inhibited tumor growth (**Figure 4J**) and reduced endpoint tumor weight in the mice model (**Figure 4K**).

**Figure 4.**
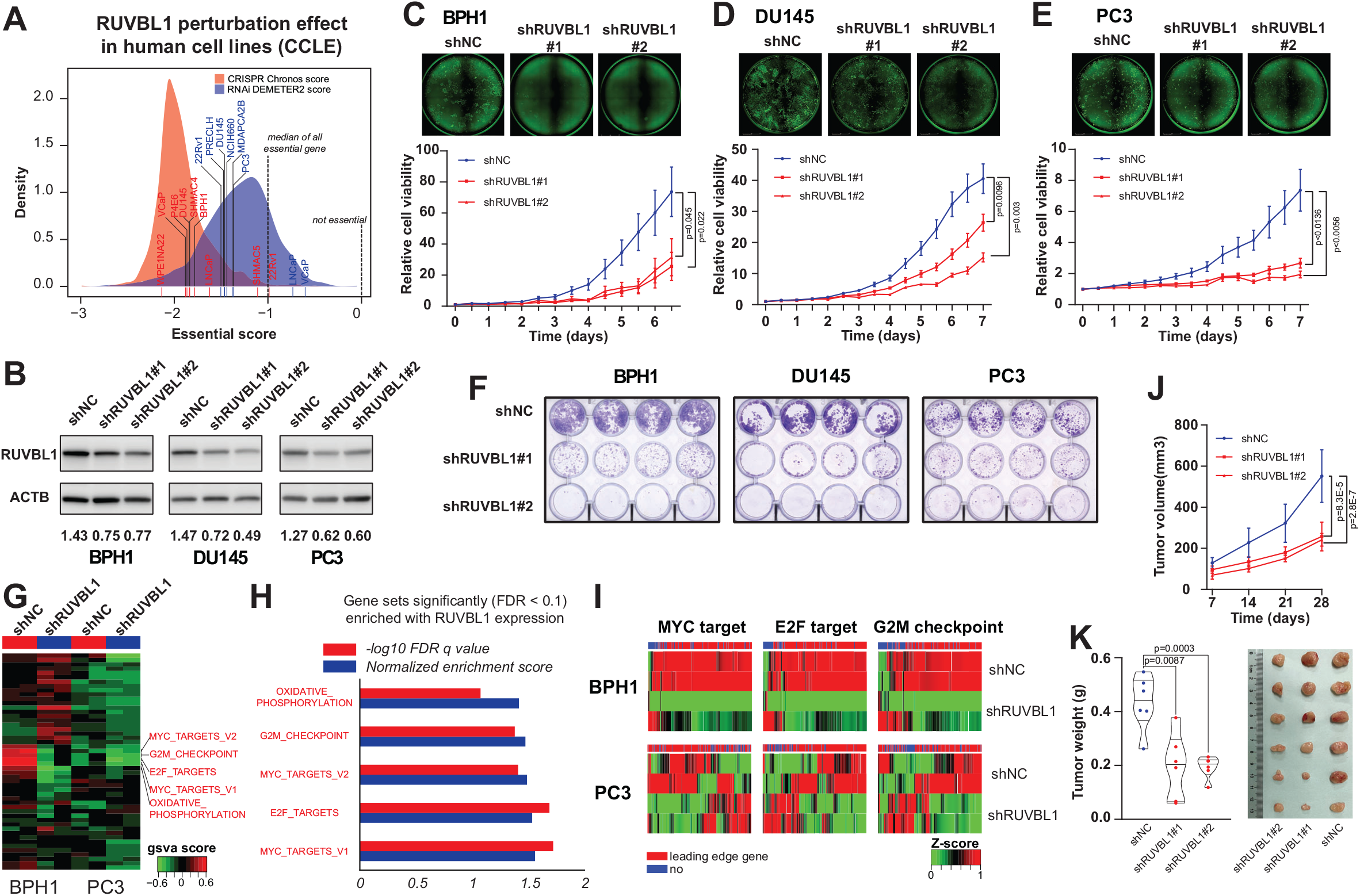
A. Dependency score of *RUVBL1* gene for CRISPR and RNAi screening from CCLE human cell line collections. A score of 0 indicates that a gene is not essential, while a score of -1 is comparable to the median of all pan-essential genes.**B.** Measuring RUVBL1 protein expression after shRNA knockdown with lentiviral infection (low MOI). **C-E.** Cell proliferation curve comparing cell growth differences after *RUVBL1* knockdown in BPH1 (**C**), DU145 (**D**) and PC3 (**E**) cell lines. **F.** Colony formation assay comparing clonogenic potential alterations after *RUVBL1* knockdown in BPH1, DU145 and PC3 cell lines. **G.** GSVA on RNA-seq data in BPH1 and PC3 cells after *RUVBL1* knockdown on HALLMARK collection pathways. **H.** GSEA summary for top rank significant *RUVBL1*-driven gene sets in BPH1 and PC3 cells. **I.** Representative heatmap showing significantly enriched HALLMARK pathways, including MYC target, E2F target and G2M checkpoint. **J-K.** Tumor volume changes (**J**) and tumor weight (**K**) comparison in xenograft experiments with PC3 cell line.

### *RUVBL1* increased aggressiveness and predicted a worse prognosis in prostate cancer

To characterize the malignant potential associated with the *RUVBL1* gene in clinical samples, we obtained three indices of genome instabilities, including the altered fraction of the genome, mutation count, and aneuploidy score for 488 TCGA prostate cancer. We found *RUVBL1* gene expression was positively associated with these indices (**Figure 5A-5C**). Additionally, we demonstrated that prostate cancer with higher *RUVBL1* expression tended to have a more advanced tumor size stage and Gleason score (**Figure 5D**). We also calculated the GSVA score for HALLMARK geneset collection with the TCGA prostate cancer RNA profiling and found that the cell cycle related pathways showed up consistently in the best 8 significantly enriched gene sets (**Figure 5E**). We further demonstrated the enrichment score(**Figure 5F-5G**) and FDR q values for these significantly enriched pathways and found consistent positive enrichment with the *RUVBL1* levels. We performed a Kaplan-Meier analysis to see whether RUVBL1 could be a prognostic marker and found significantly worse progression-free (**Figure 5H**) survivals in those with higher *RUVBL1* expression. To determine whether the *RUVBL1* enriched gene could be prognostic, we also used k-means clustering methods with the leading-edge gene to separate the prostate cancer patients into groups with different risks (**Figure 5I**). The Kaplan-Meier analysis showed that patients with increased risk tended to have significantly worse progress-free survival (**Figure 5J-5L**). Additionally, we found *RUVBL1* gene expression was consistently upregulated in PCa primary and metastasis tumor tissue (**Supplementary Figure 4B-4D**). We also demonstrated a positive association between *RUVBL1* gene expression and elevated prediagnostic PSA level, higher Gleason score, and worse biochemical recurrence-free survival in existing cohorts (**Supplementary Figure 4E-4G**).

**Figure 5.**
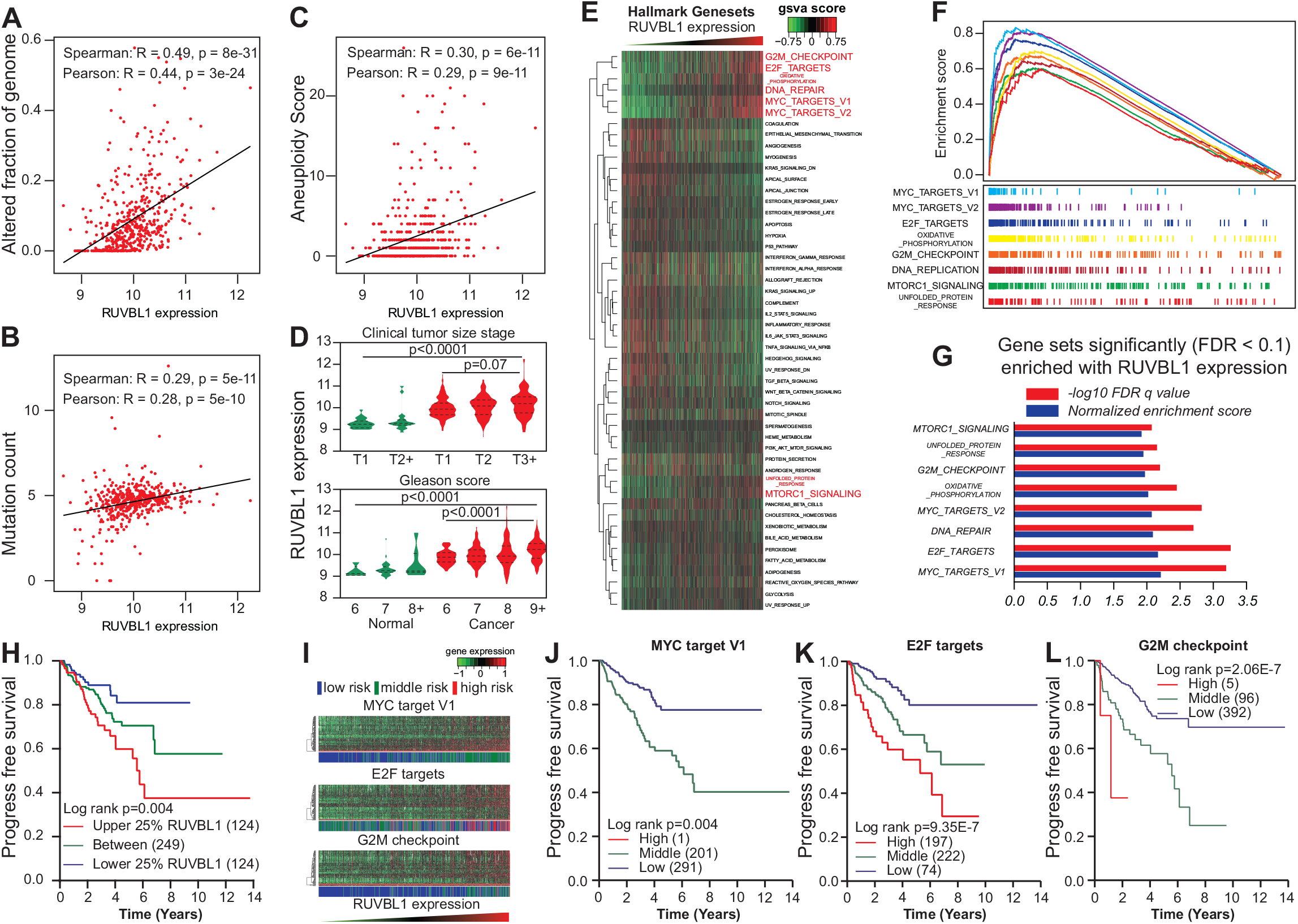
**A-C.** Associations in TCGA prostate cancer cohort between *RUVBL1* gene expression and genome instability metrics, including the altered fraction of genome (**A**), the mutation count (**B**), and the aneuploidy score (**C**). **D.** *RUVBL1* gene expression in TCGA prostate tissue with different tumor size stage and Gleason score. **E.** GSVA on RNA-seq data in TCGA prostate cancer tissue by ranking *RUVBL1* expression on HALLMARK collection pathways. **F.** GSEA enrichment plots showing significantly enriched HALLMARK pathways in TCGA prostate cohort. G. Normalized enrichment score and FDR q-value for significantly enriched HALLMARK pathways in TCGA prostate cohort. H. Kaplan-Meier analysis on progress-free survival in TCGA prostate cancer patients stratified by *RUVBL1* gene expression. **I.** Heatmap with *RUVBL1* leading-edge genes with column side bar indicating k-means stratified groups with different risks related to significantly enriched HALLMARK pathways in TCGA prostate cohort, including MYC target V1, E2F targets and G2M checkpoint. **J-L.** Kaplan-Meier analysis on progress-free survival in TCGA prostate cancer patients stratified by GSEA edge gene enriched with *RUVBL1* in each pathway, including MYC target V1 (**J**), E2F targets (**K**), and G2M checkpoint (**L**).

## Discussion

Over the past decade, GWAS and eQTL analysis have been highly productive in finding PCa risk loci and susceptible genes. ^43–45^ Despite contributing to a better understanding of the biological significance of risk predisposition, these analyses did not directly demonstrate the regulatory role of individual locus and the functional consequence of each causal gene. To better delineate the functionality of these genetic findings, a large-scale functional evaluation of target risk loci is highly warranted. Due to the non-coding nature of most risk loci, these variants are believed to play a regulatory role. Therefore, we applied the dCas9-based CRISPRi assay to target SNP sequences at risk loci, aiming to mimic the regulatory alteration caused by single base substitutions. This genome-wide screening at PCa risk loci discovered 117 SNPs showing a regulatory role in cell proliferation. Interestingly, these proliferation-related SNPs are enriched in gene transcription start sites, suggesting the majority of the phenotypical changes are related to transcription alterations caused by dCas9 interferences.

This study characterized the regulatory role of a risk SNP rs60464856. We observed consistent growth inhibition by multiple gRNAs targeting the locus. More importantly, with multiple gRNA targeting both alleles at the same genomic locations, we observed interesting phenotypical changes to seed region mismatches in real cells. As demonstrated by previous studies, ^46,47^ any mismatches at 7-bp seed region of the gRNA could cause a rapid rejection of these targets by the dCas9 protein. These facts may explain the A allele-specific depletion effect of gRNA on positions -5 to -7, as shown in our data (**see Figure 2A**). When the mismatch is located outside the seed region, the gRNA with SNP at -8 position showed depletion effects for both alleles. To further validate the regulatory role of rs60464856, we created multiple subclones carrying converted G alleles. As expected, the G allele-carrying subclones showed an elevated *RUVBL1* expression in BPH1 and DU145 cells. However, this elevated expression was not observed in PC3 subclones, possibly caused by the distinct status of histone modification and chromatin interaction in PC3 cells.

This study also showed allele-specific binding of SMC3 at rs60464856 locus, which is different from *SMC1A*. As the major subunits of human cohesin, both *SMC1A* and *SMC3* mediate multiple biological processes, including DNA looping, chromosome condensation, and chromosome segregation, by forming heterodimers. ^48,49^ Intriguingly, there have been several unique observations about the distinct phenotypical changes brought by SMC1A or SMC3 knockdown. Magdalena et al. identified that *SMC3* knockdown made *SMC1A* unstable and led to less cytoplasmic accumulation, while *SMC1A* knockdown did not influence *SMC3* stability and cytoplasmic accumulation. ^50^ A recent study^51^ reported that *SMC1A* and *SMC3* ATPase active sites had differential effects on cohesin ATPase function, and SMC3 has a unique function in DNA tethering. Our database mining showed significant signal overlap between *SMC1A* and *SMC3* binding sites in the genome, but only private peaks from *SMC3* enriched CTCF motifs in the two cell lines. As an insulator that can block enhancers to regulate target genes, CTCF was first discovered as a transcriptional repressor and believed to execute a hub role in controlling gene expression. ^42,52^ In our findings, CTCF may also play a crucial role in governing *RUVBL1* gene regulation, potentially through insulating the rs60464856 loci allelically.

*RUVBL1*, known as RuvB Like AAA ATPase 1 or TIP49, is a protein-coding gene possessing ATP-dependent DNA helicase activity and has been reported to regulate a wide range of cellular processes, ^53^ including chromatin decondensation, ^54,55^ misfolded protein aggregation, ^56^ and transcription regulation. ^57^ In addition to the previously reported mTORC1 pathway, ^58^ our enrichment analysis demonstrated that the *RUVBL1* expression was consistently correlated with cell cycle regulation and c-MYC signaling activities in both cell lines and tumor tissues. This result demonstrates a potential use of *RUVBL1* selective inhibitors in treating prostate cancer. ^59,60^ The result also suggests applying the rs60464856 genotype to stratify a target population for future clinical trials.

One potential issue with this study is inconsistent results in some tests among different cell lines. While these inconsistencies may attribute to genetic heterogeneity, we also want to highlight that some candidates found exclusively in DU145 cells are reported to be functional in prostate cells, for instance, the established functional variants residing in the binding sites of the transcription factors, TMPRSS2-ERG and HNF1B. ^61^ Additionally, to increase our gRNA library coverage, we may use novel CRISPR systems with expanded PAM site compatibility. With stringent library preparation and screening processes, the biological implications for the functional variants discovered in one cell line hits are still worth investigating, which may uncover unique SNP and gene functions specific to the cell line of interest. In summary, we applied CRISPRi screening technology to screen for survival-essential SNPs at the genome scale. We identified over a hundred functional SNPs that regulate cell proliferation. We further characterized the rs60464856 risk variant for its regulatory role in the prostate context and target gene *RUVBL1* for its functional role in prostate cancer cell proliferation and disease progression. This result will enrich our knowledge of PCa predisposition and provide insight into the cancer risk classification and potential therapeutic target for personalized treatment.

## Data and code availability

The NGS data have been archived at the Gene Expression Omnibus under the reference Series GSE224654, which includes CRISPRi screening gRNA readout (GSE224653) and RNA-seq expression for *RUVBL1* knockdown experiment (GSE224646). The gRNA sequence design, raw and normalized count, and the eQTL mapped with the candidate SNP are listed in **Supplementary Table S2**. The RIGER analysis output is listed in **Supplementary Table S3**. The publicly available datasets used are listed in **Supplementary Table S4**. The SILAC proteomics sequencing result is listed in **Supplementary Table S5**. The detailed information about the rs60464856 base editing can be accessed with the box link (https://moffitt.box.com/s/353qfjsxycaztutjufjq607hok4n3kf9).

## Acknowledgments

We thank Flow Cytometry, Molecular Genomic, Proteomic and Microscopy Core Facility at the Moffitt Cancer Center, an NCI-designated Comprehensive Cancer Center (P30CA076292). The authors also expect acknowledgment to the participants and investigators of the FinnGen study (www.finngen.fi). This work was supported by the National Institutes of Health (R01CA250018 and R01CA212097 to L.Wang., R01CA263494 to L. Wu) and National Natural Science Foundation of China (82073082) as well as the Jane and Aatos Erkko Foundation, the Finnish Cancer Foundation, and the Sigrid Juseliuksen Saatio to G-H.W. We thank Dr. Stephen N. Thibodeau at Mayo Clinic for providing the eQTL candidate list for design this screening. We also thank the PRACTICAL/ELLIPSE consortium for providing GWAS resources supporting this project. The full funding information for the consortium author is listed in Supplementary information.

## Author contribution

Y.T. and L.W. contributed to study design and performed CRISPRi and proteomics screening, statistical analysis and functional validation. D.D., Z.W. and G.-H.W. performed functional analysis and mouse work. L.Wu. and J.P. contributed to collecting GWAS summary statistics, G-H.W. and L.W. supervised the study and contributed to study design and data interpretation. Y.T., D.D., G.-H.W., and L.W. co-wrote the manuscript. All authors read, commented and approved the final version.

---

**Supplementary Figure S1:**
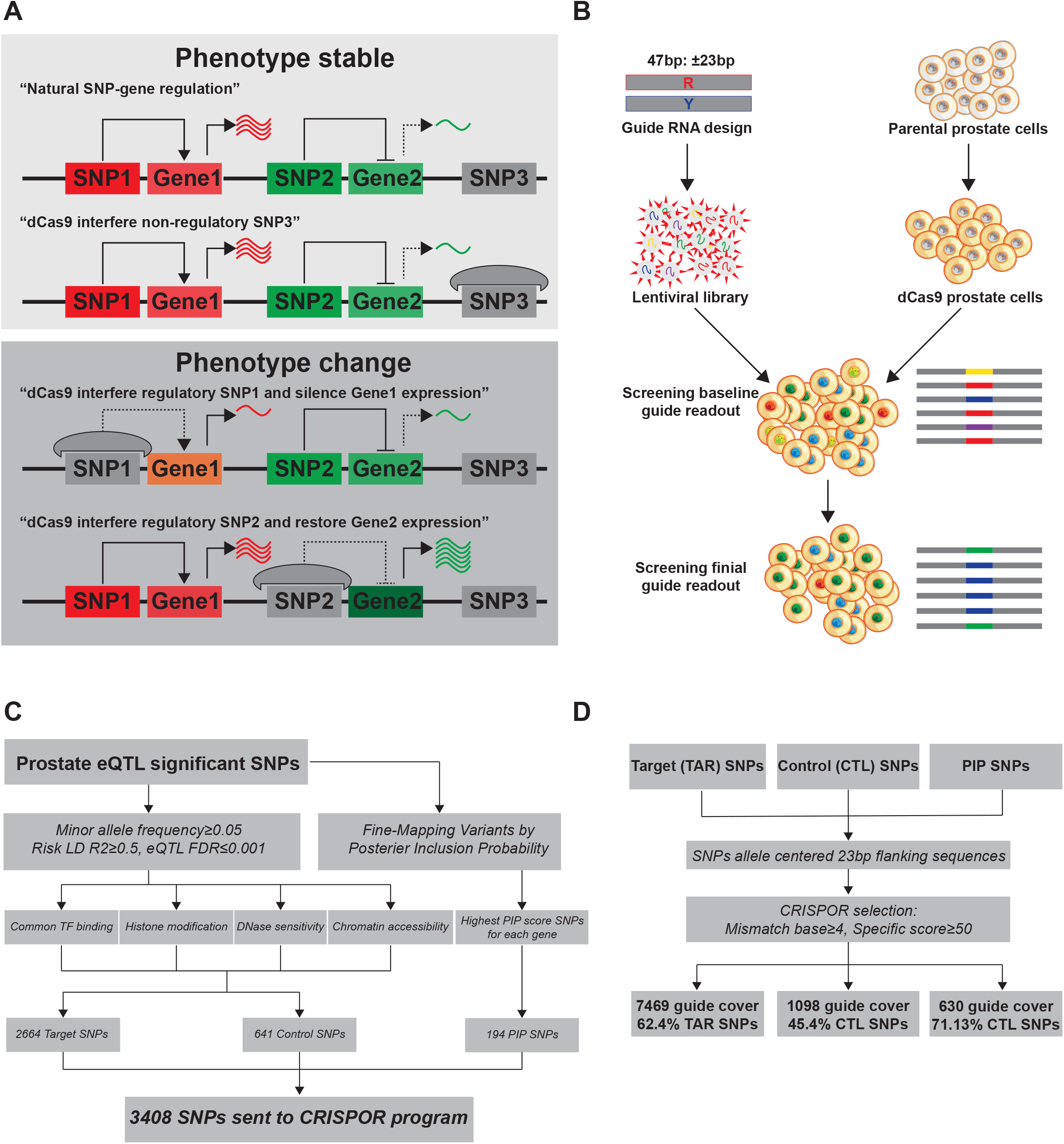
Overall CRISPRi screening projection and design. **A.** Rationale for phenotype oriented CRISPRi-SNPs-seq screening. **B.** Procedure of CRISPRi screening in dCas9 stable prostate cell lines. **C.** Candidate eQTL risk SNPs selection before gRNA design. **D.** gRNA design with CRISPOR program

**Supplementary Figure S2:**
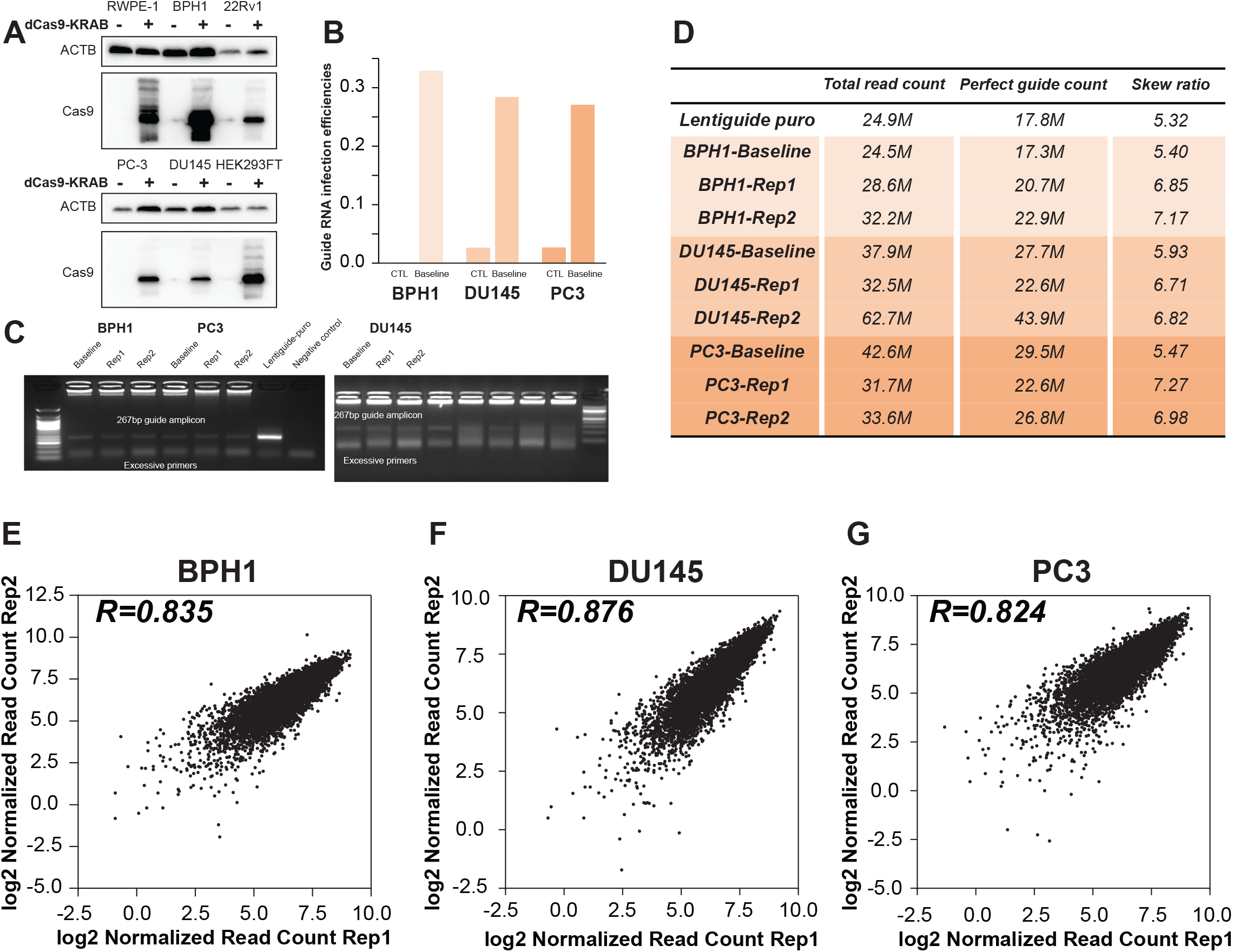
CRISPRi screening quality control. **A.** Westernblot of Cas9 protein in dCas9 stable prostate cell lines. **B.** Functional titration of lentiGuide-Puro virus to ensure low MOI integration. **C.** Gel image of gRNA readout amplicon from each screening after PCR. Template gDNA amount for each sample is equivalent to at least 5 million cells. **D.** gRNA read count summary after count_spacers.py quantification **E-G.** Reproducibility of technical replicates in BPH1 (**E**), DU145 (**F**), and PC3 (**G**) cells.

**Supplementary Figure S3:**
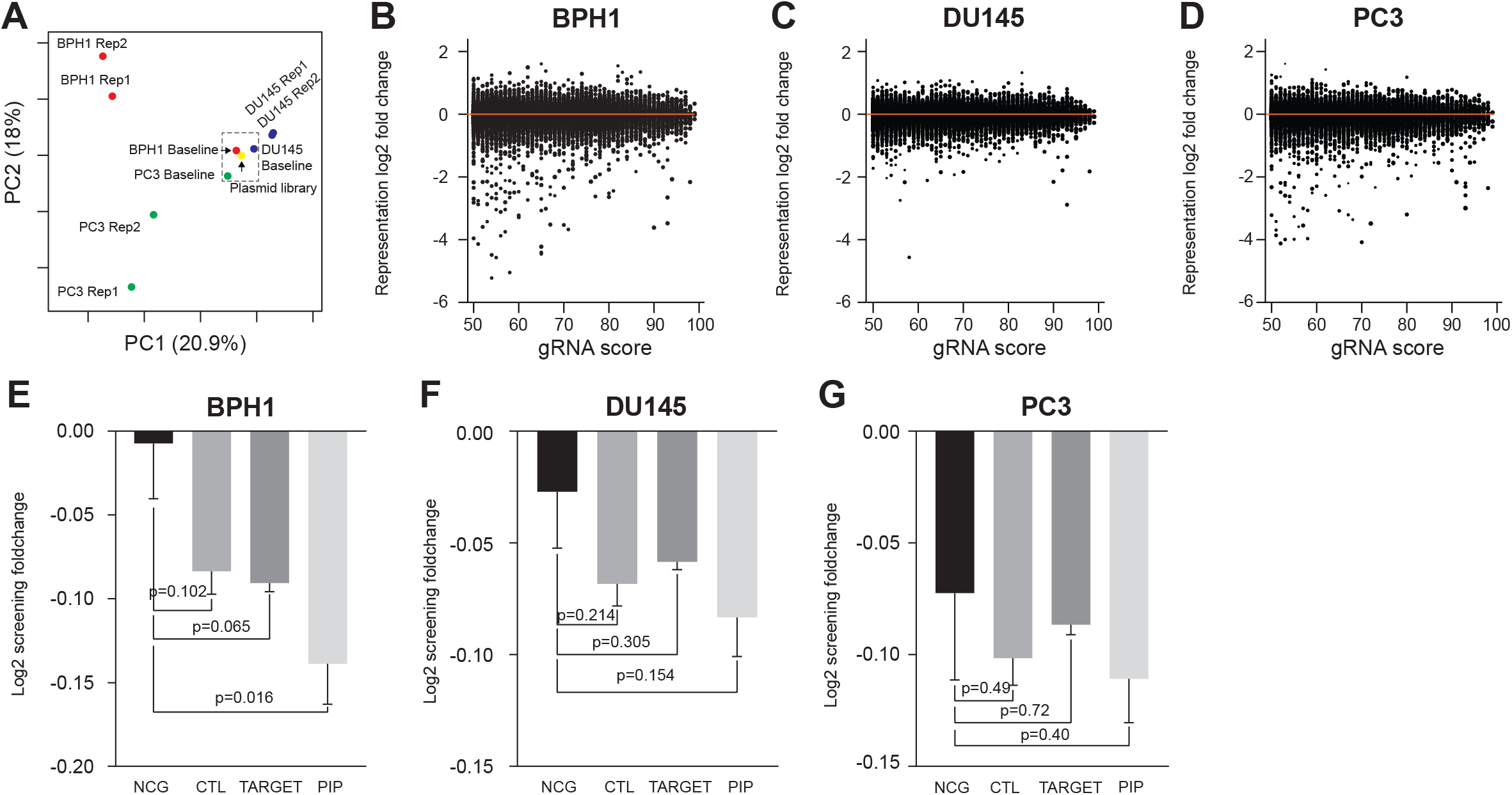
Screening candidate characterization. **A.** PCA analysis of raw gRNA count in lentiGuide-Puro plasmid and each screening sample. **B-D.** Correlation between average gRNA foldchange and specificity score in BPH1 (**B**), DU145 (**C**), and PC3 (**D**) cells. **E-G.** gRNA foldchange comparison between predefined categories in BPH1 (**E**), DU145 (**F**), and PC3 (**G**) cells.

**Supplementary Figure S4:**
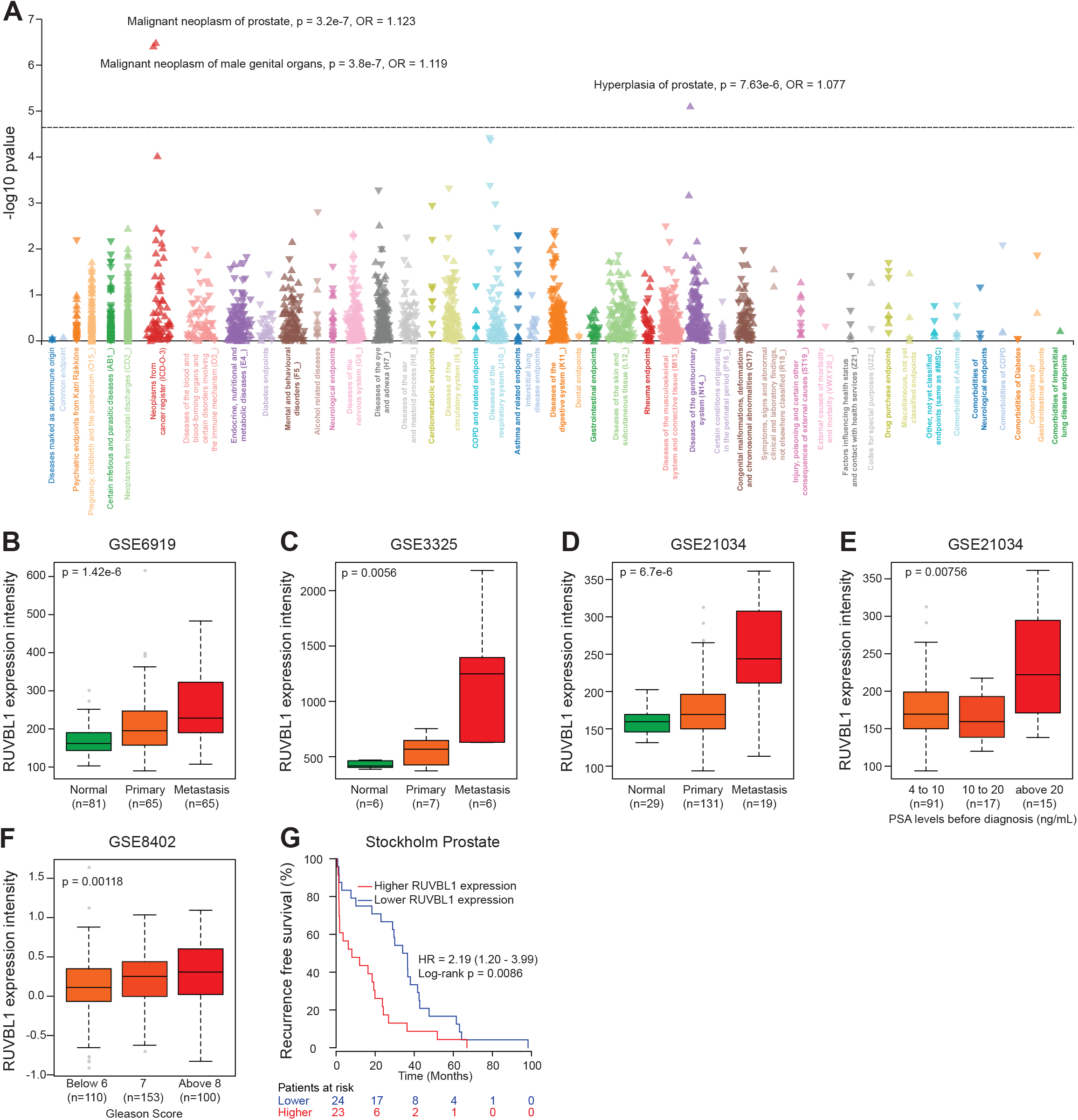
Clinical associations with RUVBL1 in other prostate cohorts. **A.** PheWAS results between rs60464856 and 2202 endpoints in the FinnGen study (n = 342,499). **B-C**. Associations between RUVBL1 gene expression and prostate cancer tissue type in Yu **(B)**, Varambally **(C)**, and Taylor’s **(D)** prostate cancer cohorts. **E.** Associations between RUVBL1 gene expression and the pre-diagnosis prostate-specific antigen (PSA) levels in Taylor’s prostate cancer cohorts. **F.** Associations between RUVBL1 gene expression and the Gleason score in Setlur’s prostate cancer cohorts. **G.** Kaplan-Meier analysis on biochemical recurrence-free survival in Stockholm prostate cohort stratified by RUVBL1 expression. The Mann-Whitney U test was used to calculate the p-value for B to F.

**Supplementary Figure S5:**
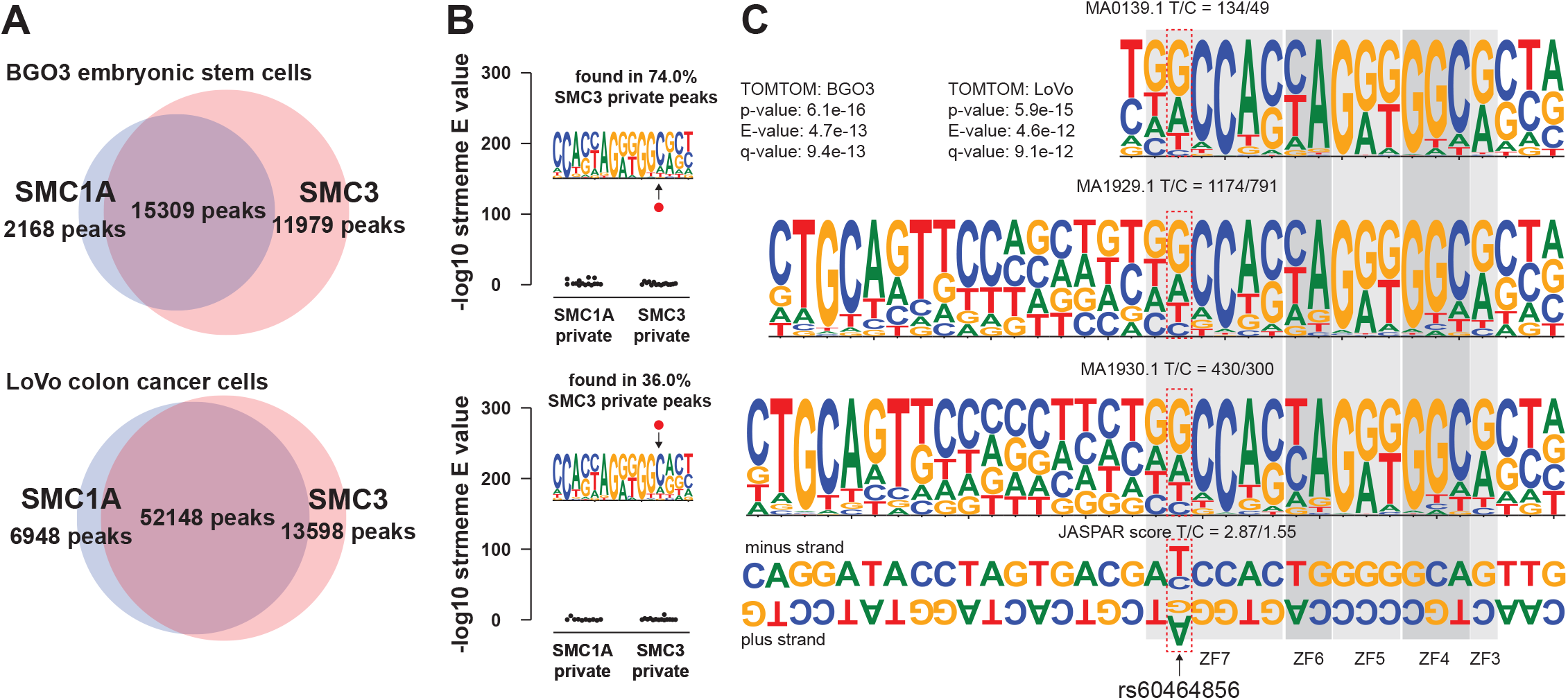
Motif analysis for SMC1A and SMC3 in human cell lines. **A.** Proportional venn diagram of SMC1A and SMC3 ChIP-seq peak overlapping in BGO3 and LoVo cell lines. **B.** STREME motif scan in BGO3 and LoVo cell lines. **C.** TOMTOM comparison of SMC3 private motif to CTCF(MA0139.1) and base composition on SNP location in different CTCF motifs. The shaded blocks highlighted DNA binding sites interacting with CTCF zinc finger (ZF) domains.

