## Supplementary material for "Combined CRISPRi and proteomics screening reveal a cohesin-CTCF-bound allele contributing to increased expression of *RUVBL1* and prostate cancer progression": Table S1

**Supplementary Table S1 Sequences of primers and oligos used in this project**

|  | Primer Sequences for QPCR |
| --- | --- |
| RUVBL1-mRNA-F | AGGTGAAGAGCACTACGAAGA |
| RUVBL1-mRNA-R | CTACTATGACGCCACATGCCT |
| ACTB-mRNA-F | CCAGAGCAAGAGAGGCATCC |
| ACTB-mRNA-R | GTACATGGCTGGGGTGTTGA |
|  | Primer Sequences for reporter assay construction |
| RUVBL1_Gibson_F | cggcggccaagcttagacacCACATCTCACGTTGCAAG |
| RUVBL1_Gibson_R | aacagtaccggattgccaagTCTTCATTTTGCAGACGC |
|  | shRNA sequences |
| shRUVBL1#1-Sense | GUGGCGUCAUAGUAGAAUUAA |
| shRUVBL1#1-Antisense | UUAAUUCUACUAUGACGCCAC |
| shRUVBL1#2-Sense | CCGGCCAACUUGCUUGCUAAA |
| shRUVBL1#2-Antisense | UUUAGCAAGCAAGUUGGCCGG |
| shNC-Sense | CCUAAGGUUAAGUCGCCCUCG |
| shNC-Antisense | CGAGGGCGACUUAACCUUAGG |
|  | Primers for genome editing |
| rs60464856-A2G-guide | TGGATCGTCACTAGGTATCC |
| rs60464856-Sanger-F | TAATTCCCGCTGTATCCCAGTGTC |
| rs60464856-Sanger-R | CCCGCCATTATTTCCTCAGGGAAGT |
| ARMS-F-outer | AACCGTCCCATAGCCTGCCACTGCATTC |
| ARMS-R-outer | AGAGGTGTGGCCAGTGGACCAGGGAGTT |
| ARMS-R-inner | GGGGCCGCCCCAGGATACCTAGTGACTAC |
|  | Primers for ChIP-qPCR |
| rs60464856-locus-F | TAATTCCCGCTGTATCCCAGTGTC |
| rs60464856-locus-R | CCCGCCATTATTTCCTCAGGGAAGT |
| rs60464856-NC-F | AAGTGAGGCATTCTATGGGACTG |
| rs60464856-NC-R | CCAGGGGATATTCCTCTGTGC |
| AS-rs60464856-R-A | CCAGGATACCTAGTGACGAC |
| AS-rs60464856-R-G | CCAGGATACCTAGTGACGAT |
| AS-rs60464856-F | CAAAGCCCTGCAGTAACTAACC |
