## Supplementary material for "Combined CRISPRi and proteomics screening reveal a cohesin-CTCF-bound allele contributing to increased expression of *RUVBL1* and prostate cancer progression": Table S4

**Supplementary Table S4 Published datasets used in this paper**

|  | Accession number and web link |
| --- | --- |
| DU145 – H3K4me1 | SRR3624829, SRR3624830 |
| DU145 – H3K4me3 | SRR3624831 |
| DU145 – H3K27ac | SRR5823947 |
| PC-3 – H3K4me1 | ENCSR566UMF |
| PC-3 – H3K4me3 | ENCSR275NCH |
| PC-3 – H3K27ac | ENCSR826UTD |
| PrEC – H3K4me1 | SRR1282226 |
| PrEC – H3K4me3 | SRR1282227 |
| PrEC – H3K27ac | SRR1282224 |
| RWPE-1 – H3K4me1 | SRR1645120, SRR1645121 |
| RWPE-1 – H3K4me3 | SRR1645122, SRR1645123 |
| RWPE-1 – H3K27ac | SRR1645108, SRR1645109, SRR1645110, SRR1645111 |
| BGO-SMC1 | SRR445918 |
| BGO-SMC3  LoVo-SMC1 | SRR445917  SRR952473 |
| LoVo-SMC3 | SRR952474 |
| Yu’s PCa cohort | GSE6919 |
| Varambally’s PCa cohort | GSE3325 |
| Taylor’s PCa cohort | GSE21034 |
| Setlur’s PCa cohort | GSE8402 |
| Stockholm’s PCa cohort | GSE70769 |
